# Decreased utilization of allocentric coordinates during reaching movement in individuals with autism spectrum disorder

**DOI:** 10.1101/2020.07.15.204016

**Authors:** Yumi Umesawa, Takeshi Atsumi, Reiko Fukatsu, Masakazu Ide

## Abstract

Despite numerous reports of abnormalities in limb motor controls in spatial orientation in individuals with autism spectrum disorder (ASD), the underlying mechanisms have not been elucidated. We studied the influence of allocentric coordinates on ongoing reaching movements, which has been reported to strongly affect the reaching movements of typically developing (TD) individuals. ASD and TD participants observed a target presented randomly on one of the four corners of a frame on a screen. After it disappeared, another frame was presented slightly shifted leftward/rightward. The participants touched the memorized position of the target relatively congruent with a reference frame (allocentric condition) or ignoring it (egocentric condition). Results suggested that touch positions were less affected by shift directions of reference frame in the allocentric condition in ASD participants, so that they tended to reach in egocentric manner. Our findings demonstrate that decreased utilization of visual landmarks in ongoing movement may underlie motor disabilities in autism.

## Introduction

Individuals with autism spectrum disorders (ASD) often show poorer fine motor skills such as handwriting, drawing and ball handling (1). Another study pointed out that handwriting difficulties in autistic children might be resulting from impairments in visuo-motor transformation (2). While visual information of target object is useful in acquiring internal model for adaptive motor controls in typically developing (TD) children (3), children with ASD tended to put greater reliance on proprioceptive feedbacks in motor learning (Haswell et al., 2009). This character was found to relate to reduced synaptic volume especially in the cerebellum which is a representative neural basis of internal models of motor control (4). From these findings, different optimizing strategies of motor control in internal model would be responsible for difficulty to learn adaptive movements.

However, it is more essential to immediately and abruptly approach some objects in our daily life, before acquiring strategies for adequate movements. In these immediate movements, not only recognition of the object itself but also recognition of it in relation to surrounding environment of the object, that is, allocentric coordinates, is useful for efficient goal-directed reaching (5, 6). Egocentric coordinates are another set of clues for reaching by locating the target in relation to the body, head, retina and so on. A previous study showed that the reaching for targets were more accurate and less variable when the target location provided a context for allocentric coordinates (7–11). It has been demonstrated that background coordinates (i.e., reference frame: RF) were also essential in the situation of immediate saccade and reach toward a target (12, 13).

While a large body of evidence for difficulties of limb control required visuo-motor transformation in individuals with ASD (14–17), it still has not been verified that abnormal characteristics in utilizing allocentric coordination during reaching movements. The purpose of this study is to clarify whether individuals with ASD can be affected by allocentric coordinates in an immediate goal-directed ongoing reaching, which did not need motor learning.

## Materials and Methods

### Participants

We recruited 17 individuals with ASD and 17 TD individuals (Table 1). ASD participants were recruited from parent groups for individuals with developmental disorders and the Department of Child Psychiatry in the Hospital of National Rehabilitation Center for Persons with Disabilities. There was no significant between-group difference in age and handedness, which was assessed using the laterality quotient score of the Edinburgh Handedness Inventory (18). The participants also completed the Japanese version of the Autism spectrum quotient (AQ) scale (19, 20), in which higher scores indicate stronger autistic traits. AQ scores in ASD group were significantly higher than those of the TD group (two-tailed *t*-test: *t* (32) = 4.02, *p* = 0.0003, Cohen’s *d* = 1.38, 95% CI = [5.55, 16.31]). The Intelligence Quotients (IQs) were assessed using the Wechsler Adult Intelligence Scale-Third Edition (WAIS-III). There were no significant between-group differences in Verbal IQ (VIQ), performance IQ (PIQ) and FIQ. The present study was approved by Ethics committee of the National Rehabilitation Center for Persons with Disabilities, and all participants gave informed written consent before the experiment.

**Table 1.**
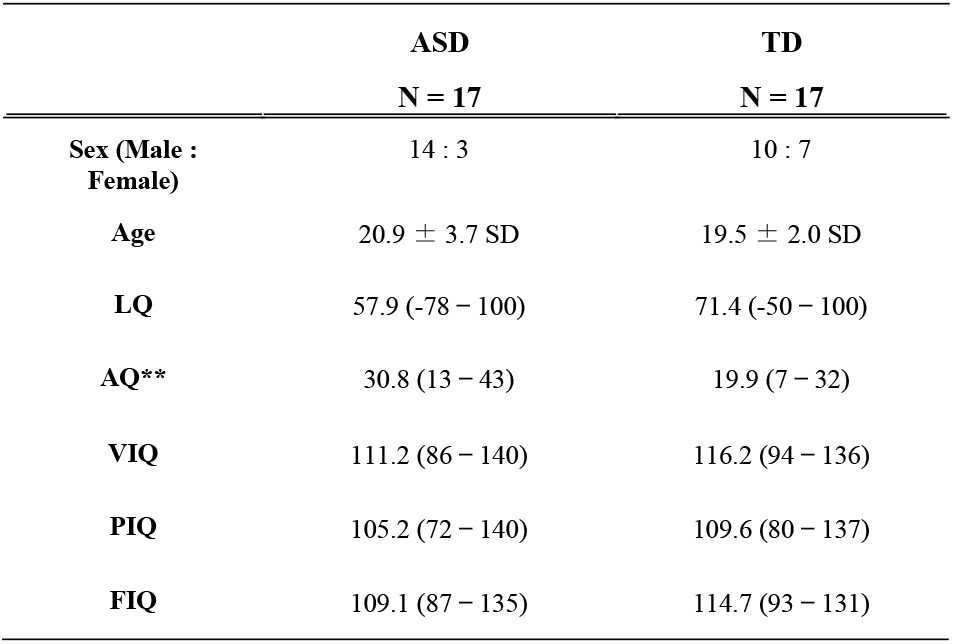
Information of participants. Scores in each cell and parentheses denote the averages and ranges in ASD and TD groups, respectively. Asterisks denote statistical significance (\*\**p* < 0.01).

| Sex (Male : Female) | 14 : 3 | 10 : 7 |
| --- | --- | --- |
| Age | 20.9 ± 3.7 SD | 19.5 ± 2.0 SD |
| LQ | 57.9 (-78 – 100) | 71.4 (-50 – 100) |
| AQ** | 30.8 (13 – 43) | 19.9 (7 – 32) |
| VIQ | 111.2 (86 – 140) | 116.2 (94 – 136) |
| PIQ | 105.2 (72 – 140) | 109.6 (80 – 137) |
| FIQ | 109.1 (87 – 135) | 114.7 (93 – 131) |

### Apparatus and general task procedures

According to the experimental paradigm in a previous study (13), the participants performed reaching movements for a target position in relation to a reference frame (RF) on a screen. The participants were seated facing a touch screen (ET1903LM-2UWA-1-WH-G, dimensions 376.32 mm (H) × 301.06 mm (V), Elo Touch Solutions, Inc.) with a resolution of 1280 pixels ×1024 pixels placed 360 mm in front of their eyes with their heads restrained by a chin rest (see Fig 1A). A customized PC (Predator G5900, Acer Inc.) and PsychoPy3 (21) were used to control the experiment. An index finger stand was positioned 200 mm below and 100 mm ahead of a subject’s eyes in the midsagittal plane. A target (a yellow cross, 15 mm = 2.4° in visual angle) and a RF (Standard RF; a white square, 80 mm = 12.7°) appeared on a black background for 200 ms after a random delay (1800 – 2000 ms) followed by a blank (500 ms). The target cross was presented at a random location on either of the four corners of the frame. Another reference frame (Test RF) was displayed after 500 ms blank and shifted 20 mm (3.2°) either leftward or rightward in the horizonal plane compared with the position where the Standard RF was presented. The participants touched the position on the screen with their index finger soon after the Test RF was displayed according either to instructions: “Touch the disappeared target position ignoring Test RF” (*Egocentric condition*), or, “Touch the disappeared target position in Test RF relatively congruent with the position in Standard RF” (*Allocentric condition*). They were instructed to touch the target location as quickly and precisely as possible without any intentional corrections. They rested the finger on the finger stand within 500 ms after touch. It took 4 – 5 s to complete one trial (Fig 1B). Each of the *Egocentric* and *Allocentric conditions* included 60 trials, and it took 5 – 6 min to complete. The participants performed reaching tasks in both conditions on different days to avoid the influence of the effect of learning a strategy that is efficient for a specific condition (i.e., *Egocentric* or *Allocentric*). The orders of the conditions were counter-balanced among participants.

**Fig 1.**
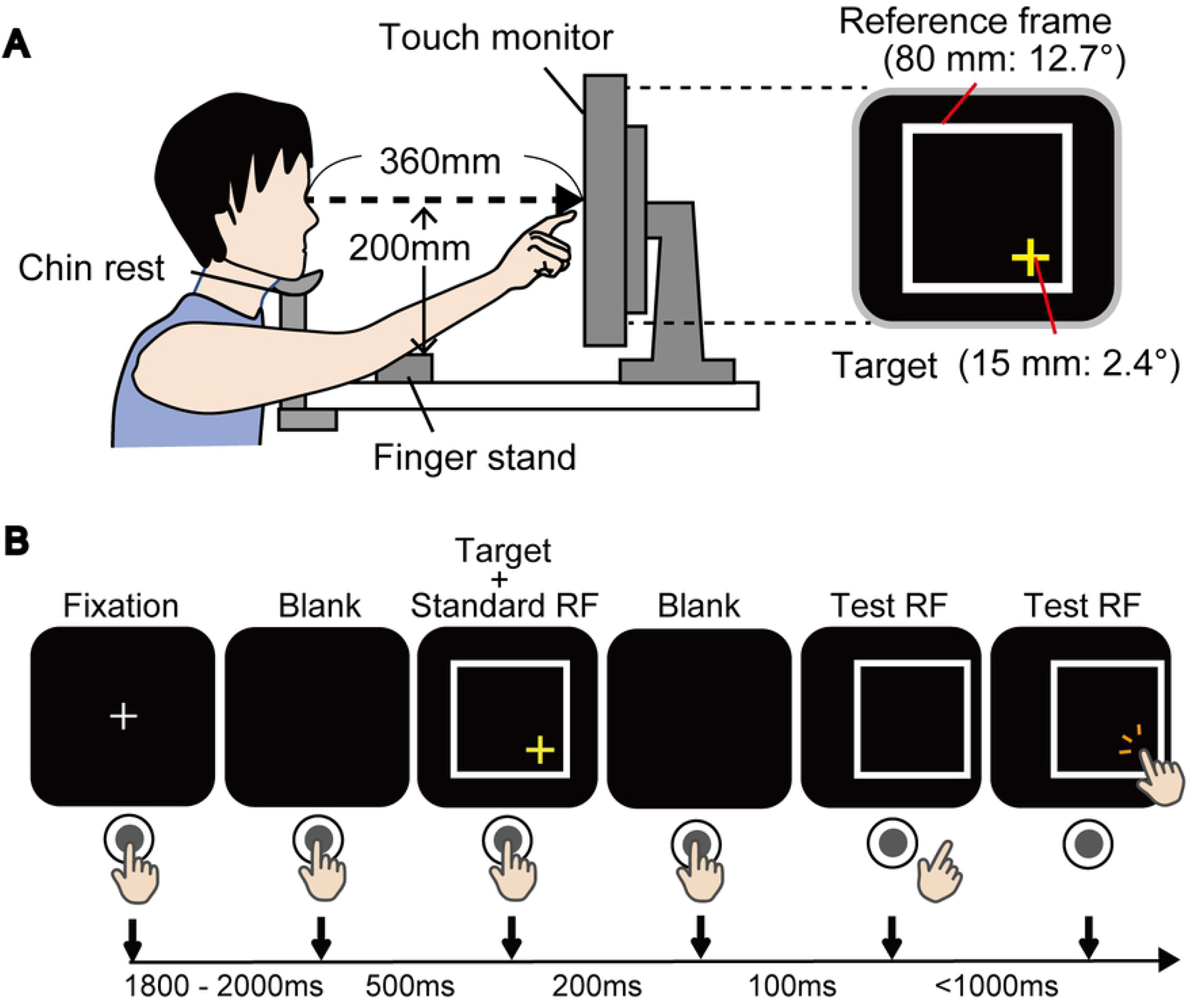
Experimental set up. Experimental set up. **(A)** Experimental setups. Participants were seated facing a screen placed 360 mm in front of their eyes with their heads restrained by a chin rest. A yellow cross target was presented randomly on either of the four corners within a white square (i.e., reference frame: RF) on the screen. **(B)** Task procedures. Each column is the schematics of the visual field and the position of the hand relative to the button and the screen. A target with the Standard RF appeared for 200 ms proceeded by a blank of 500 ms and a fixation of random duration between 1800 – 2000 ms. After a blank (100 ms), only a Test RF shifted either leftward or rightward. The participants touched a position on the screen according to instructions (see Methods section) as quickly as possible (within 1000 ms). Test RF disappeared when the participants touched the screen.

### Evaluation indexes

Response times were calculated as time elapsed from the presentation of the Test RF to the timing of touch on the screen. We calculated the amount of errors in X-axis to evaluate influence of allocentric representation in the Egocentric condition (AR errors) and that of egocentric representation in the Allocentric condition (ER errors) on the touch position using following formula (see Fig 2).

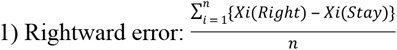

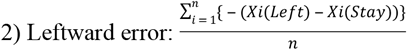

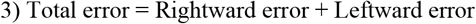

1) 2) Rightward and Leftward errors in *Egocentric (or Allocentric) condition* were estimated as the degree how touch position in Test RF translated (or did not translate) toward memorized target position relatively congruent with Standard RF. 3) Total error (AR error / ER error) was the summation of both the errors.

**Fig 2.**
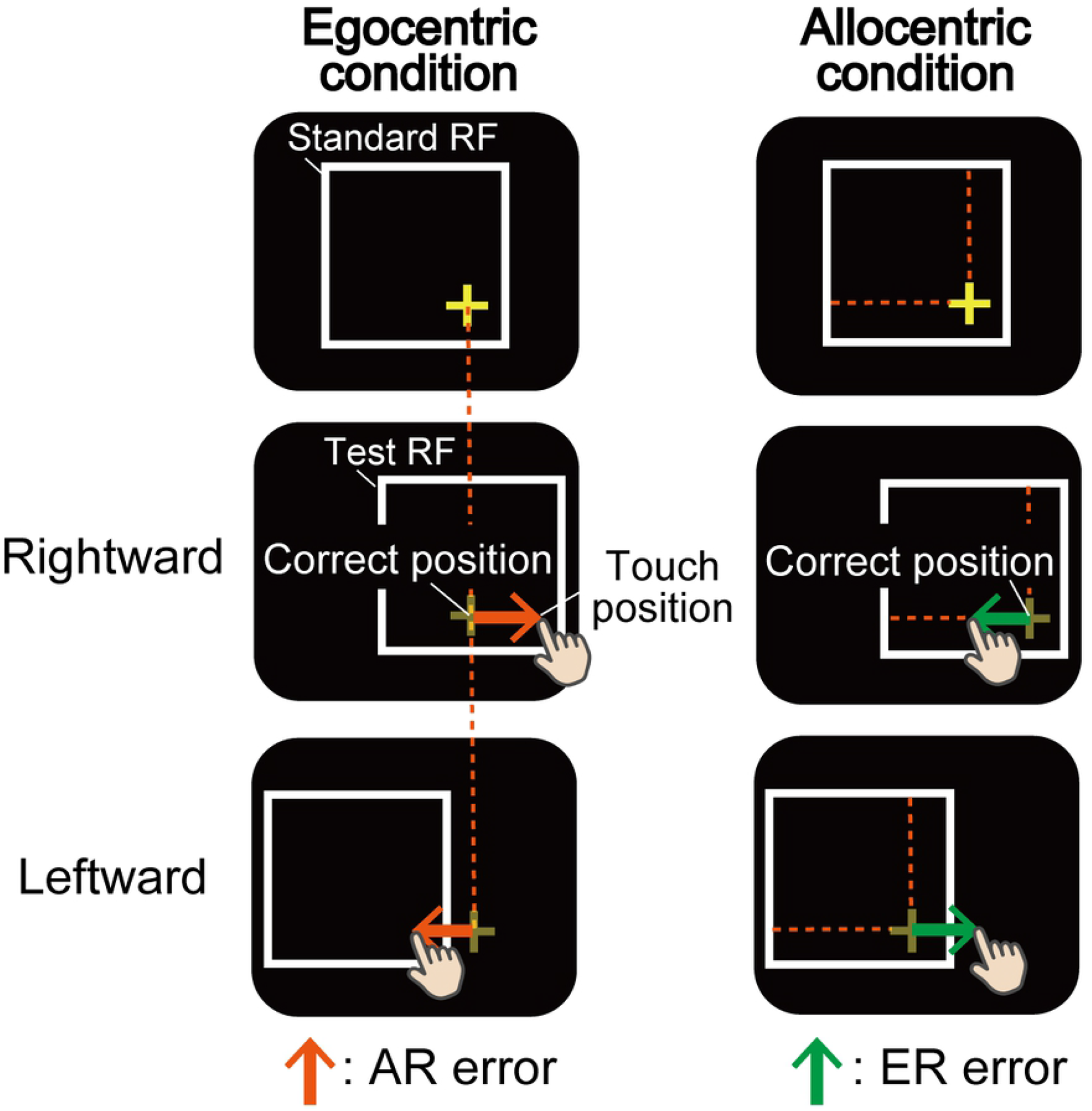
Evaluation indexes. We calculated the amount of error in X-axis to evaluate the influence of each coordinate’s conditions (i.e., *Egocentric* or *Allocentric conditions*) on the touch position. In the *Egocentric condition*, orange arrows indicate the amount of translation of the touch affected by allocentric representation (i.e., the target position in Standard RF) (AR errors). In the *Allocentric condition*, green arrows indicate the amount of translation of touch ‘*not*’ affected by allocentric representation, but rather affected by egocentric representation (ER errors).

### Statistical Methods

We used data that were obtained within 1 sec from the start of the measurement of responses for subsequent analysis. We performed one-sampled *t* test versus zero in AR errors and ER errors to examine whether the translated touch position was affected by the allocentric or egocentric representations in the *Egocentric* or *Allocentric conditions*, respectively. Student *t* tests was used for comparing group’s means as for AR or ER errors. We calculated Cohen’s *d* to demonstrate effect size of the difference of the groups. We used SPSS Version 23.0 (IBM, New York, U.S.) to perform *t* test, and G*power 3.1 (22) to calculate the effect sizes.

## Results

Fig 3A and 3B depict touch positions in each condition. By calculating average of X-axis errors of touch position from correct position for each participant, we estimated AR errors in the *Egocentric condition* and ER errors in the *Allocentric condition* (Fig 3C).

**Fig 3.**
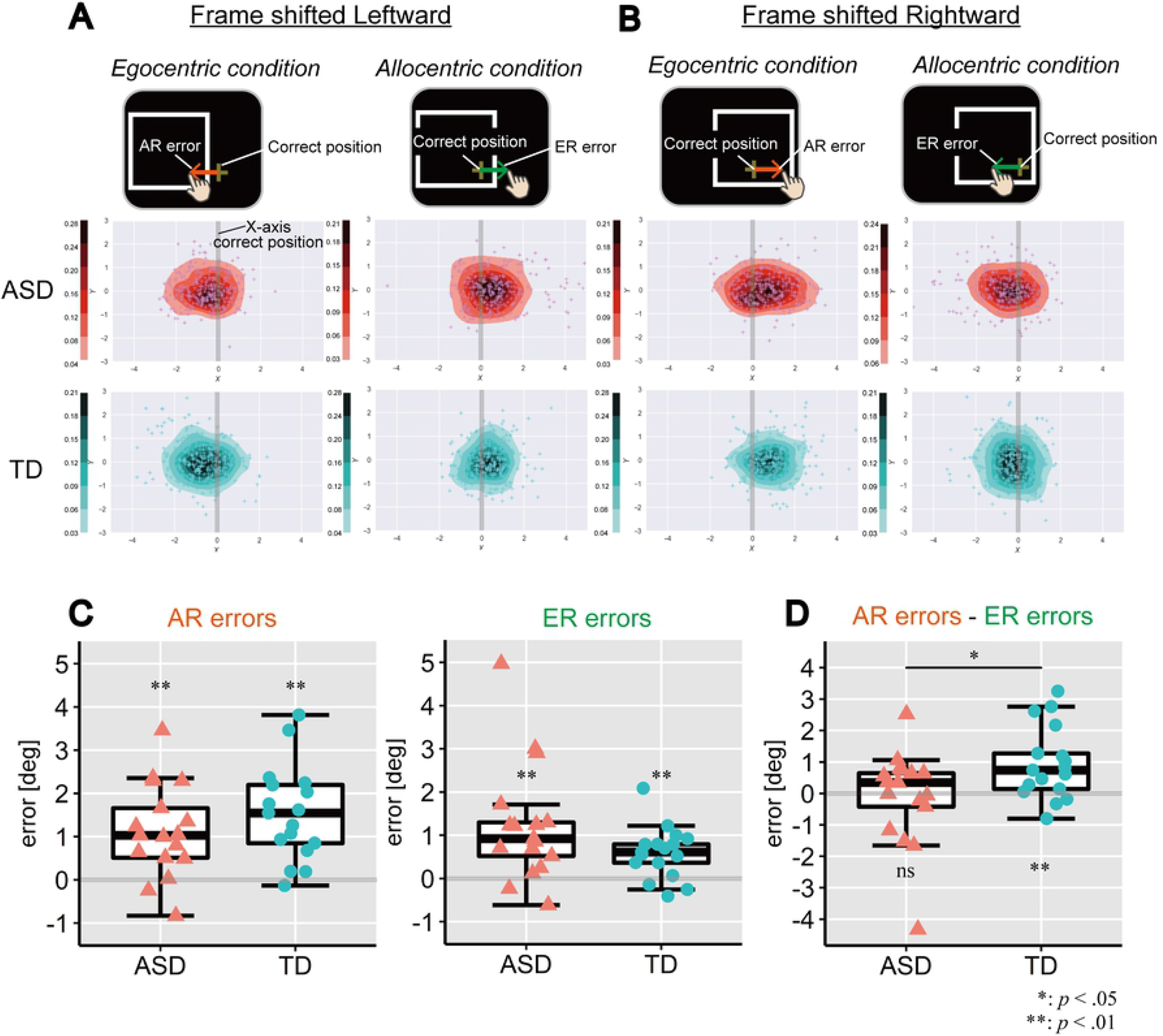
Behavioural results in ASD and TD groups. **(A)** Touch positions on the screen when frame was shifted leftward. X- and Y-axes indicates horizontal and vertical dimensions (deg) of the screen. Depth of colour (red: ASD, blue: TD) of the heat maps depict the proportions of frequent touches in the area estimated by probability density function. Grey vertical lines indicate correct positions of touch according to instructions (allocentric or egocentric) in X-axis. **(B)** Touch positions on the screen when frame was shifted rightward. Others are same as above (A). **(C)** Group comparisons of touch errors in each condition. Left box plot denotes AR errors in the *Egocentric condition*. Right box plot denotes ER errors in the *Allocentric condition*. The upper and lower boundaries of the standard boxplots are at the 25^th^ and 75^th^ percentiles. The horizontal line across the box marks the median of the distribution. The ends of vertical lines below and above the box represent the minimum and maximum, respectively. **(D)** The amount of preferential use of allocentric representation by subtracting ER errors from AR errors.

Both AR and ER errors were significantly greater than zero each in the *Egocentric* and *Allocentric conditions* in both ASD and TD groups (AR errors: *t*s (16, 16) = [4.37, 5.80], *ps* = [<0.01, <0.01], *d*s = [1.06, 1.41], 95% CIs = [{0.58, 1.67}, {0.97, 2.10}], in ASD and TD groups, respectively; ER errors: *t*s (16, 16) = [3.74, 4.08], *p*s = [<0.01, < 0.01], *d*s = [0.91, 0.99], 95% CIs = [{0.53, 1.92}, {0.28, 0.8}], in ASD and TD groups, respectively). There were no significant group difference in AR errors in the *Egocentric condition (t* (32) = −1.11, *p* = 0.28, *d* = 0.38, 95% CI = [−1.16, 0.34]), while ER errors in the *Allocentric condition* in ASD group was slightly greater than TD group (*t* (32) = 1.78, *p* = 0.085, *d* = 0.61, 95% CI = [−0.09, 1.37]).

We calculated the amount of preferential use of allocentric representation by subtracting ER errors from AR errors within each participant. Positive values indicate how the participant’s touch were affected by shifted reference frame. Results demonstrated that TD participants significantly preferred to use allocentric representation (one-sample *t* test from zero: *t* (16) = 3.4, *p* = 0.004, *d* = 0.82, 95% CI = [0.35, 1.54]), while ASD participants did not show such a preference for using allocentric representation (*t* (16) = −0.27, *p* = 0.79, *d* = 0.07, 95% CI = [−0.86, 0.66]). Group comparison suggested that ASD group less preferred to use allocentric representation compared with TD group (*t* (32) = −2.3, *p* = 0.029, *d* = 0.79, 95% CI = [−1.97, −0.12]) (Fig 3D).

## Discussion

We examined to what extent individuals with ASD depend on visual information of surrounded environment in feedback-absent reaching behaviour. Results demonstrated that both individuals with ASD and TD were affected by directions of shifted reference frame in ongoing reaching for the target. Although TD individuals were apt to reach in allocentric manner rather than egocentric manner, ASDs did not show this prioritize: ASDs had difficulty in spatially orienting their arm by referring to the presence of Test frame contrary to TDs. Our present findings provide a new insight in terms of motor difficulties in ASD with respect to the utilization of allocentric coordination in immediate ongoing movement before acquiring the internal model (Haswell et al., 2009; Izawa et al., 2012; Marko et al., 2015).

The TD participants in the present task likely stored the target location relative to allocentric coordinates to decode the spatial position whereas the ASD participants depended more on egocentric coordinates in visually guided reaching behavior. While allocentric spatial representation is available by 2 years of age (23), incomplete maturation of allocentric representation were observed in children with TD between 6-7 years old (24). Since allocentric coordinates is useful for precise limb movement (7–11), individuals with TD come to preferentially employ the allocentric coordinates compared to egocentric coordinates in a developmental stage. This development of utilization of allocentric coordinates may be immature in ASD.

Allocentric spatial representation is thought to be processed in the ventral stream of the brain projecting from the primary visual cortex (V1) to the inferior occipito-temporal cortex, while dorsal stream projecting from V1 to the posterior parietal cortex (PPC) is assumed to process egocentric representation (25). Recent functional magnetic resonance imaging (fMRI) studies have suggested that the occipito-temporal network associate with the representation of allocentric coordinates (26–28). A review of 48 diffusion tensor imaging (DTI) studies found that ASD brain tended to have decreased white matter tracts that pass through the temporal lobe (29). Previous literatures for neural basis of motor disabilities in ASD frequently reported functional and/or structural abnormality in the action observation network (30), involving several frontal regions to inferior parietal lobule (31, 32). We speculate that lower connectivity in occipito-temporal network would associate with the reduced utilization of allocentric coordinates for movement in spatial navigation.

We found that individuals with ASD utilize allocentric coordinates less compared with TD participants in ongoing reaching movement, and rely on egocentric coordinates relatively strongly. These characteristics observed in every reach movement may be a basis of difficulty for acquiring effective internal model using visual information for motor control in learning, and contribute to difficulties in visually guided motor function in daily life.

## Acknowledgements

We would like to thank M. Chakrabarty for the valuable comments in constructing the experiment. We are also grateful K. Matsushima for advices from a view point of occupational therapist. We appreciate Y. Wang for technical helps in taking assessments.

